# Modulating Transthyretin Fibril Stability with D-Retro-Inverso Peptides

**DOI:** 10.64898/2026.08.12.744519

**Authors:** Lucy M. Coleman, Ulrich H. E. Hansmann

**Affiliations:** Department of Chemistry & Biochemistry, University of Oklahoma, Norman, Oklahoma 73019, United States

## Abstract

A major cause of heart failure in elderly patients are deposits of Transthyretin (TTR) fibrils. Using molecular dynamic simulations, we explore how the stability of TTR fibrils can be modulated by D-Retro-Inverso (DRI) Peptides, built from D-amino acids with the sequence of the parent peptide switched, and describe a mechanism by which one of these peptides, DRI-K6V, disrupts TTR fibrils. Our results may open the way to design of peptide drugs targeting established TTR amyloidosis.

## INTRODUCTION

The extracellular protein Transthyretin (TTR) assembles as a tetramer transporting thyroxin and retinol throughout the body. Upon dissociation of the tetramer, the released monomers can partially unfold and aggregate into insoluble fibrils that may damage heart and nervous system but can also affect eye, kidney, and the gastrointestinal tract.^1^ Wildtype transthyretin amyloidosis (ATTRwt), where amyloid deposits are found mainly in the heart, is estimated to affect over 300,000 elderly patients worldwide, while the hereditary (i.e., mutant-based) form of disease, transthyretin amyloidosis (ATTRv), is seen in about 50,000 patients.^2–4^ Early detection of the disease is rare and the mean survival times in untreated patients are about 2.5-3.6 years after diagnosis.^5–8^ Current treatment options are limited to management and preventing aggravation of symptoms. Heart transplants can replace the diseased hearts in patients with ATTRwt or severe ATTRv. Liver transplants can remove in patients with ATTRv the source of mutant TTR,^3^ and RNAi treatments, patisiran and vutrisiran, can inhibit expression of TTR.^9–12^ Other U.S. Food and Drug Administration (FDA)-approved drugs, such as tafamadis,^13^ diflusinal,^14, 15^ and acoramidis,^16, 17^ prevent aggregation into fibrils by stabilizing the TTR tetramer. Common to all these approaches is that they reduce the progression of fibril formation but cannot eliminate the existing and disease-symptoms causing amyloid deposits. Hence, as existing therapies have limited effectiveness during late stages of a disease, there is a need for exploring avenues to inhibit or even reverse TTR fibril formation.

One possibility is the use of small peptides that can bind to TTR fibrils, destabilizing and dissolving them. In the context of other amyloid forming human proteins, short peptides have been shown to inhibit aggregation by acting as β-sheet breakers.^18, 19,20–22^ Well studied are peptides that are built from the proteolysis-resistant D-amino acids as they avoid short *in vivo* lifetimes that would limit their therapeutic use.^23^ One possibility are D-retro-inverso (DRI)-peptides, where reverting the amino acid sequence and flipping the chirality of amino acids from L-enantiomers to D-enantiomers leads to molecules that resemble the parent peptide and often maintain their biological activity while avoiding proteolysis.^24^ For instance, we showed in previous work that *in silico* a five-residue peptide, RSFFS, and its DRI form, DRI-R5S, destabilizes serum amyloid A (SAA) fibrils.^25^ A subsequent *in vivo* study demonstrated that in a mice model treatment with DRI- R5S after myocardial infarction decreases formation of SAA fibrils resulting in improved heart function and recovery.^26^ Motivated by these previous results, we propose a computational framework using virtual screening to identify and evaluate DRI peptides for their ability to destabilize TTR amyloid fibrils, and molecular dynamics (MD) simulations to identify their mechanism of action. While previous studies focused on short peptides that inhibit fibril formation and growth by interacting with the TTR monomer,^27^ binding to soluble oligomers or capping fibril ends, ^28–32^ we are interested in peptides that can disrupt mature fibrils potentially opening treatment strategies for patients who have already developed amyloid deposits.

We remark that we did not address the specificity of our peptides candidates, i.e., their binding to competing proteins in the heart or other organs. This is because we are mainly interested in understanding the mechanism by which peptides can destabilize TTR fibrils, a question difficult to access with current experimental approaches. To our knowledge there are only two (transgenic double-humanized) mice strains (hV/hV:hR/hR and hV/hM:hR/hR) where TTR amyloids are observed in the heart, and that therefore could serve as a mice model.^33^ However, amyloid deposition is seen only at 24 month of age, i.e. in mice of advanced age, and appears to be very sensitive to environmental conditions. This makes *in vivo* experiments of TTR amyloidosis difficult and costly. Hence, a detailed knowledge of mechanism by that peptides can dissolve the TTR fibril is needed for optimizing potency and selectivity of potential drug candidates prior to experimental validation. For this purpose we identify in the present paper suitable peptides and try to understand the mechanism by that they cause destabilization of the TTR fibril. Using high-throughput docking and Rosetta energy analysis to screen a library of candidate peptides, we select final candidates s based on predicted engagement with dynamically vulnerable regions of the fibril as identified by MD simulations of an experimentally resolved TTR fibril model. MD simulations of the top candidates interacting with TTR fibrils are used to validate promising binders and allow us to identify the mechanism by which one of these peptides, DRI-K6V, disrupts TTR fibrils.

## METHODS

### Generation of Fibril Model

The TTR fibril model was generated using the cryo-EM derived structure of a patient-derived wildtype TTR fibril at a resolution of 2.78 Å as deposited in the Protein Data Bank (PDB) with PDB ID: 8ADE.^34^ Several other studies have also posed models of TTR fibrils, all revealing a common fold with several unresolved regions: the first ten residues of the N-terminus (residues G1-C10), the last four residues of the C-terminus (residues N124-E127), and a 21-residue segment in the middle of the protein (residues A36-H56).^35–37^ For our simulations we added the missing N- and C-terminal regions using the MODELLER software for de novo loop modeling.^38^ On the other hand, the unresolved segment (residues A36-H56) was not added under the hypothesis that it is proteolytically cleaved during fibril formation, consistent with mass spectrometry data showing that most fibril chain fragments begin or end within this region.^36, 37, 39^ Hence, our fibril model of five TTR chains consists two "proto fibrils" made of fragments G1-K35 and G57-E127, see **Figure 1a**.

**Figure 1.**
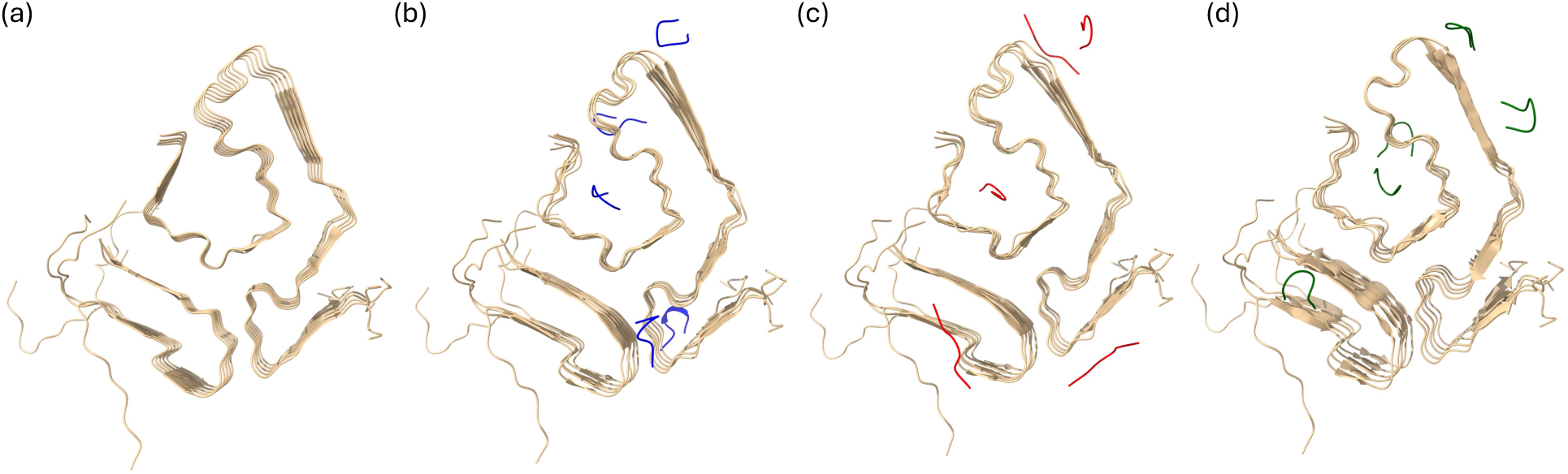
Start conformations of simulations of (a) a sole TTR fibril model (our control), and in presence of three DRI-peptides: (b) DRI-K6V (in blue), (c) DRI-W6I (in red), and (d) DRI-W6Y (in green).

### Peptide Library Generation

Amyloidogenic regions of the TTR protein sequence were identified using web-based software TANGO^40^, AGGRESCAN^41^, ZipperDB^42^, WALTZ^43^ and PASTA^44^. We focus on hexapeptides based on extensive studies on six-residue amyloidogenic peptides, which is around the length of a typical β-strand.^45, 46^ All unique six-residue sequences were extracted from amyloidogenic regions of TTR using a sliding window approach. The library was generated through mutational analysis of the six-residue amyloidogenic sequences where amino acids were replaced only with such belonging to the same group as defined by the chemical properties of their side chains (**Table ST1**). We considered only one-point and two-point mutated peptides, i.e., such that differ from the parent sequence by only one or two residues. In this way, we generated a total of 9,297 peptides.

The solubility of each of these peptide was evaluated by the Kyte-Doolittle Hydropathy^47^ scale, and the β-strandness of each peptide was screened by the Chou and Fasman^48^ parameters. Peptides with moderate hydrophobicity (0.8-1.8) and moderate overall β-strand propensity (1.0-1.2) were considered for further screening to ensure structural stability for target binding, reduced tendency to aggregate, and solubility, all characteristics seen in inhibitory peptides targeting other amyloid diseases.^49, 50^ Next, the peptides were screened for amyloidogenicity by AmyloGram^51^, AGGRESCAN^41^, and ZipperDB^42^. Sequences identified to be amyloidogenic were used for molecular docking, generating a final library of 1,134 peptides. In addition to our library of peptides based on amyloidogenic sequences of the TTR protein, we also considered the WALTZ-DB 2.0 database^46^ containing 1,404 unique hexapeptides with amyloid forming potential.

All considered peptides were converted to their DRI form by reversing the sequence, generating structures with PeptideBuilder^52^, and converting each amino acid from the L-enantiomer to the D- enantiomer by swapping the positions of the β-carbon and α-hydrogen. The resulting peptides were energy minimized implicit solvent using SANDER from AmberTools with the Generalized Born implicit solvent model.^53^

### Peptide Library Screening

For an initial high-throughput docking we used AutoDock Vina version 1.2.5 with a grid search area of 81.810 Å x 95.640 Å x 58.440 Å and center coordinates of 60.565, 60.070, 60.560 Å.^54^ Peptides were categorized as one-point mutations, two-point mutations, or WALTZ peptides. The top 100 scoring peptides from each category were selected for further analysis.

Based on the AutoDock Vina binding poses, we observed that the peptides preferred docking to the central polar cavity. Additionally, our 400K molecular dynamics simulations of the fibril identified residues K80-S100 as an early region to destabilize. These residues are part of the top beta loop and include side chains (e.g. Phe, Leu, Trp, Tyr, Ile) forming fibril-stabilizing interchain contacts. Therefore, we refined our search to peptides that could bind to the central cavity and perturb this hotspot region. Starting from the top 100 AutoDock Vina binding poses of each category (one-point mutations, two-point mutations, and WALTZ peptides), all complexes were subjected to constrained FastRelax in Rosetta, and the lowest-energy model was retained. Binding energetics were evaluated using Rosetta InterfaceAnalyzer to compute the binding free energy (dG_separated), buried surface area (dSASA_int), interfacial hydrogen bonds (hbonds_int), and buried polar atoms (delta_unsatHbonds) (**Table ST2**). Binding free energy estimates ΔG are reported in Rosetta Energy Units (REU). Per-residue energy decomposition was performed for the top 10 candidates, extracting both one-body and pairwise terms.

Candidates were analyzed by their Rosetta binding free energy estimate, contacts with hotspot residues K80-S100 per chain (weighted by per-residue fluctuation from the 400K control simulation, see discussion in the Results section), hydrophobicity, and per-residue energy decomposition at the hotspot region. The five candidates showing the strongest perturbation of residues K80-S100 (DRI-KALGIV, DRI-YTIGAL, DRI-WTFNLY, DRI-VALHGF, DRI- WKALLI) were selected for independent validation with HADDOCK 2.4^55^ using residues of the fibril cavity as active residues. The resulting structures clustered by root-mean-square-deviations (RMSD) between each other, and the lowest-energy structure from the top-ranked cluster was selected for downstream analysis. HADDOCK poses were relaxed and scored using the same Rosetta protocol. Final candidates were selected based on per-residue energetics at the hotspot region across both AutoDock-Rosetta and HADDOCK-Rosetta analyses. The candidates were also prioritized for potential mechanistic diversity, as their per-residue energy patterns suggested distinct modes of cavity engagement. Three peptides were selected as final candidates for MD simulation–DRI-KALGIV (DRI-K6V), DRI-WKALLI (DRI-W6I), and DRI-WTFNLY (DRI- W6Y). DRI-K6V and DRI-W6I are one-point mutations generated from amyloidogenic sequences of TTR, and DRI-W6Y comes from the WALTZ-DB 2.0 database. DRI-K6V and DRI-W6I are derived from adjacent segments of TTR ^80^KALGIS^85^ and ^79^WKALGI^84^, respectively. In the native TTR protein, the ^80^KALGIS^85^ segment spans the C-terminal end of the EF helix (T75-L82) and the start of the EF loop (G83-E89), a region that stabilizes both the strong and weak dimer interfaces in the homotetramer but becomes conformationally dynamic at acidic pH, with hereditary mutations in this region accelerating aggregation.^56, 57^ Additionally, this segment has been identified through crystallography as an amyloidogenic core of TTR, forming an in-register steric zipper as in the fibril spine.^58^ DRI-K6V preserves the native Leu82/Ile84 hydrophobic pair that forms the steric zipper, with the substitution of Ser to Val adding a branched aliphatic side chain that increases β-strand propensity and hydrophobicity.^58^ DRI-W6I has a Trp residue capable of both hydrophobic packing and π- stacking and replaces the native Gly with Leu increasing in this way the overall hydrophobic surface area. DRI-W6Y, an amyloidogenic peptide from the WALTZ-DB database, is derived from residues 140-145 of KCNIP1, a calcium-binding EF-hand protein that regulates Kv4-type voltage-gated potassium channels.^46^ It contains large aromatic residues (Trp, Phe, Try) which support both hydrophobic contacts and π-stacking interactions.

We remark that our three peptide candidates form upon docking compact conformations that permit insertion into the polar cavity of the TTR fibril, potentially disrupting the interstrand hydrogen bonding networks, electrostatic interactions, and hydrophobic packing that maintain the fibril, see **Figures 1b-1d**. On the other hand, the TabF2 and TabH2 TTR-targeting peptides previously developed by the Eisenberg group^59^ adopt upon docking an extended conformation stretched on the surface perpendicular to the fibril axis, see **Supporting Information Figure SF1**. As this arrangement is effective preventing further growth of fibrils, but does not destabilize existing fibrils, we did not consider TabF2 and TabH2 in our study.

### General Simulation Protocol

To generate the initial structures for the molecular dynamic simulations, all starting structures were converted to GROMACS format using pdb2gmx, and hydrogens were added to the structures. The N- and C-terminals of the fibril were left charged, and the ends of the resolved fragments were also left charged as if they were cleaved. The N- and C-termini of the peptides were capped with an NH_2_ and CONH_2_ group, respectively, in order to counteract electrostatic repulsion between the peptide’s terminal groups. The fibril contains five chains, and the peptides were docked to the surface of the fibril in a 1:1 ratio using HADDOCK 2.4 software.^55^ For each system, the first peptide was docked to the central polar cavity, while the remaining four were docked using random surface-exposed patches. Sketches of the so-generated start conformations are shown in **Figure 1**, and the corresponding atomic coordinates are given as **Supporting Information** (**TTR-fibril- coordinates.zip**). Note that the peptides were free to move or disengage with the fibril throughout the simulation.

Depending on availability of computational resources, simulations utilized either the GROMACS 2022 or 2020.4 software:^60^ GROMACS 2022 for the control and DRI-K6V systems and GROMACS 2020.4 for DRI-W6I and DRI-W6Y systems. All simulations relied on the CHARMM36m all-atom force field^61^ and TIP3P explicit water model,^62^ and identical simulation parameters and analysis protocols were applied across all runs. The fibril system was placed in a cubic box with a minimum distance of 15 Å between the solute and the edge of the box and periodic boundary conditions. The system was solvated and counter ions were added to neutralize the system in accordance with the particle-mesh Ewald method. Na^+^ and Cl^-^ ions were added at physiological concentration of 150 mM. The system was energy minimized for up to 50,000 steps using the steepest descent method. The system then underwent a NVT equilibration at constant volume and temperature of 310K for 200 ps, followed by a 200 ps NPT equilibration at a constant pressure of 1 atm. Heavy atoms were constrained with a force constant of 1000 kJ/mol•nm^2^. Production runs were performed at a constant temperature of 310K and constant pressure of 1 atm. We considered three independent trajectories of 500 ns each to capture global structural changes in the presence of the peptides.

The temperature of the simulations was maintained using a velocity-rescale thermostat^63^ with a coupling constant of 0.1 ps, and pressure was controlled with the Parrinello-Rahman barostat^64^ with a relaxation time of 2 ps. Equations of motion were integrated with a 2 fs timestep, enabled by constraining water geometry with the SETTLE^65^ algorithm and protein bonds containing hydrogen atoms to their equilibrium lengths with the LINCS^66^ algorithm. Periodic boundary conditions were applied to all systems, with long-range electrostatics computed using particle-mesh Ewald with a real-space cutoff of 12 Å and a Fourier grid spacing of 1.6 Å. Short-range van der Waals interactions were truncated at 12 Å with smoothing from 10 Å. Each system was simulated in triplicate with differing initial velocity distributions. The total number of atoms and water molecules from each system as well as the length of trajectories for each system is listed in **Table 1**, and the corresponding atomic coordinates of the start and final conformations are given as **Supporting Information** (**TTR-fibril-coordinates.zip**).

**Table 1.** Simulated Systems

| System | Total Atoms | Water Molecules | Individual Simulation Time (ns) | Total Simulation Time (ns) |
| --- | --- | --- | --- | --- |
| Control | 176,260 | 56,037 | 500 | 1500 |
| DRI-K6V | 191,683 | 61,016 | 500 | 1500 |
| DRI-W6I | 216,748 | 69,338 | 500 | 1500 |
| DRI-W6Y | 197,847 | 63,039 | 500 | 1500 |

### Trajectory Analysis

MD trajectories were analyzed with the GROMACS tools^60^ and visualized using VMD^67^ and UCSF ChimeraX^68^. GROMACS tools were used to calculate root-mean-square deviation (RMSD), root-mean-square fluctuation (RMSF), radius of gyration (R_g_), solvent-accessible surface area (SASA) with a spherical probe of 1.4 Å, and pairwise distances used for contact analysis. All contacts are defined at a cutoff of 4.5 Å between heavy atoms in a residue pair. Stacking contacts refers to contacts between chains of the fibril while packing contacts are defined as contacts between the N-terminal fragment (residues G1-K35) and the C-terminal fragment (residues G57- E127).

## RESULTS AND DISCUSSION

In order to evaluate the ability of our three peptides to destabilize and disrupt the TTR fibril, we compare molecular dynamics simulations where the peptides interact with the fibrils with such where the peptides are absent. The evolution of the fibril conformations was followed for each system in three independent trajectories over 500 ns at physiological temperature of 310K and standard pressure of 1 atm. The atomic coordinates of the final conformations of all trajectories are also given as Supporting Information in the compressed folder **TTR-fibril-coordinates.zip**. To quantify structural changes, we have calculated as a function of time the root-mean-square deviation (RMSD) over all heavy atoms with respect to the start conformation. This quantity changes little with time in the control simulations and in all cases approaches within 300 ns a plateau, see **Figure 2**. Hence, when calculating averages, we consider only the last 200 ns of our trajectories. A visual inspection of the final conformations (see the insets to **Figure 2**) shows little changes, confirming the known stability of the TTR fibril, with structural changes mainly in the polar cavity of residues G57-G83. This is not surprisingly as residues A36-H56 are assumed to be proteolytically cleaved in during fibril formation, i.e., our fibril model consists of two "proto- fibrils", and we expect flexibility of the end residues K35 and G57.

**Figure 2.**
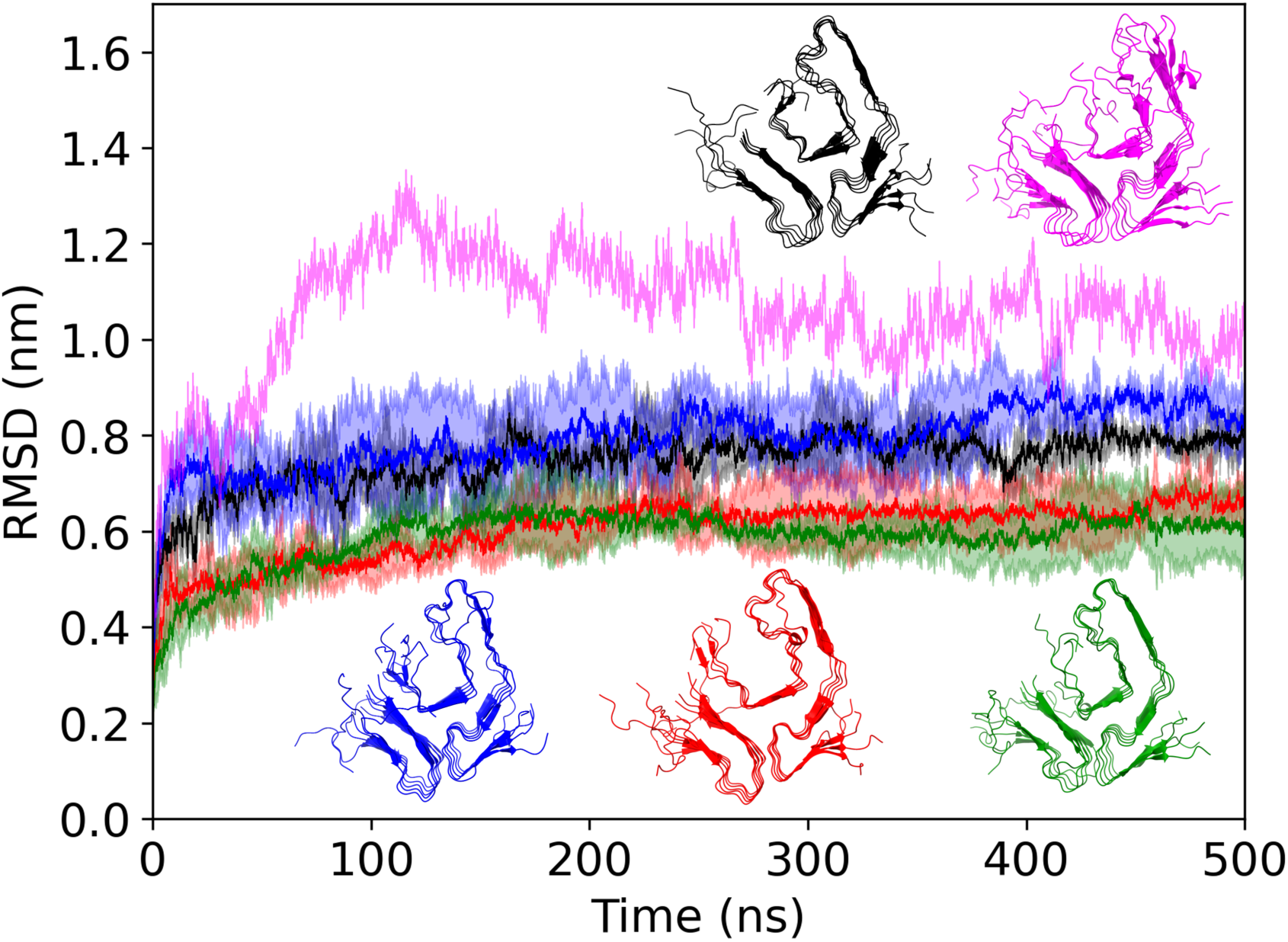
Root-mean-square deviation (RMSD) to the start configuration, calculated over all heavy atoms, as function of time for the TTR fibril in presence of DRI-K6V (blue), DRI-W6I (red), DRI-W6Y(green), and the control (black). We draw the RMSD for the additional control simulations at the unphysiological high temperature of 400K in magenta. The final conformations of all five systems are drawn using the same color coding.

This increased flexibility for residues G57 to G83 is also seen in **Figure 3** where we show the residue-wise root-mean-square fluctuation (RMSF) calculated over the last 200 ns and averaged over all five chains. However, interestingly we see in **Figure 3** a slightly raised RMSF for the segment of residues K80-S100, corresponding to the loop connecting strand G (residues K76-K80) of the polar cavity with the strand H (residues V93-N98), indicating flexibility in the spatial arrangement between the cavity and the adjacent strand H. However, this signal is weak, and to obtain a clearer picture we added another control simulation at an unphysiologically high temperature of 400K where we expect a weakening of the fibril. The RMSD as function of time and the final conformation of this trajectory are also shown in **Figure 2**, both indicating a partial dissociation of the TTR fibril, with not only the polar cavity but also the strand H distorted. Comparing the residues-wise RMSF of this trajectory with the ones from simulations at the physiological temperature of 310K we see now a clear signal for increased flexibility of the K80- S100 segment. This flexibility allows in the biologically active structure of the TTR protein (which assembles into a homotetramer) hydrogen bonding between the segments of residues A91-A97 on adjacent strands of the four chains in the dimer of dimers. Hence, we hypothesize that this flexibility, while needed in the native structure, leads to a structural weakness in the fibril that is of little relevance at physiological temperatures but becomes noticeable at raised temperature. It therefore seems likely that a peptide inhibitor would also act on this region, and this hypothesis was a guiding assumption in the search for binding sites for the peptides to the TTR fibril.

**Figure 3.**
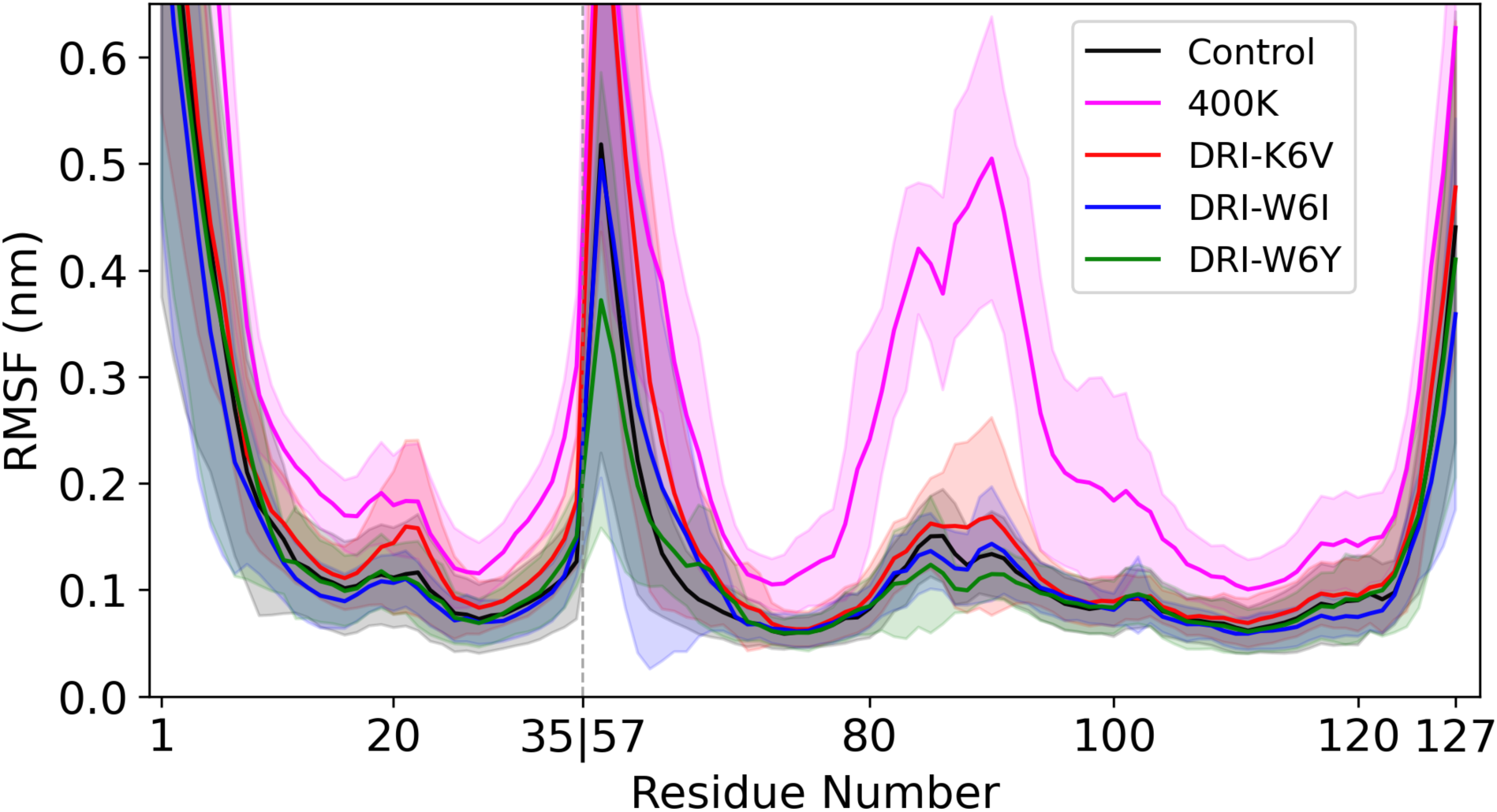
Residue-wise root-mean-square fluctuation (RMSF) calculated for each system over the last 200 ns and averaged over all three trajectories and five chains in the TTR fibril.

**Figure 1** shows the start conformation of three sets of simulations where our DRI-peptides in a one-to-one ratio are interacting with the TTR fibril. In all three cases are at least some of the peptides binding with the polar cavity in the initial conformation. However, these binding sites are not fixed, and the peptides move over the course of a trajectory. We show in **Figure 4a-4c** the probabilities of the three peptides to bind to a specific TTR residue as measured over the last 200 ns and averaged over three independent runs. Corresponding binding probabilities for the peptide residues are shown in **Figure 5a-5c**.

**Figure 4.**
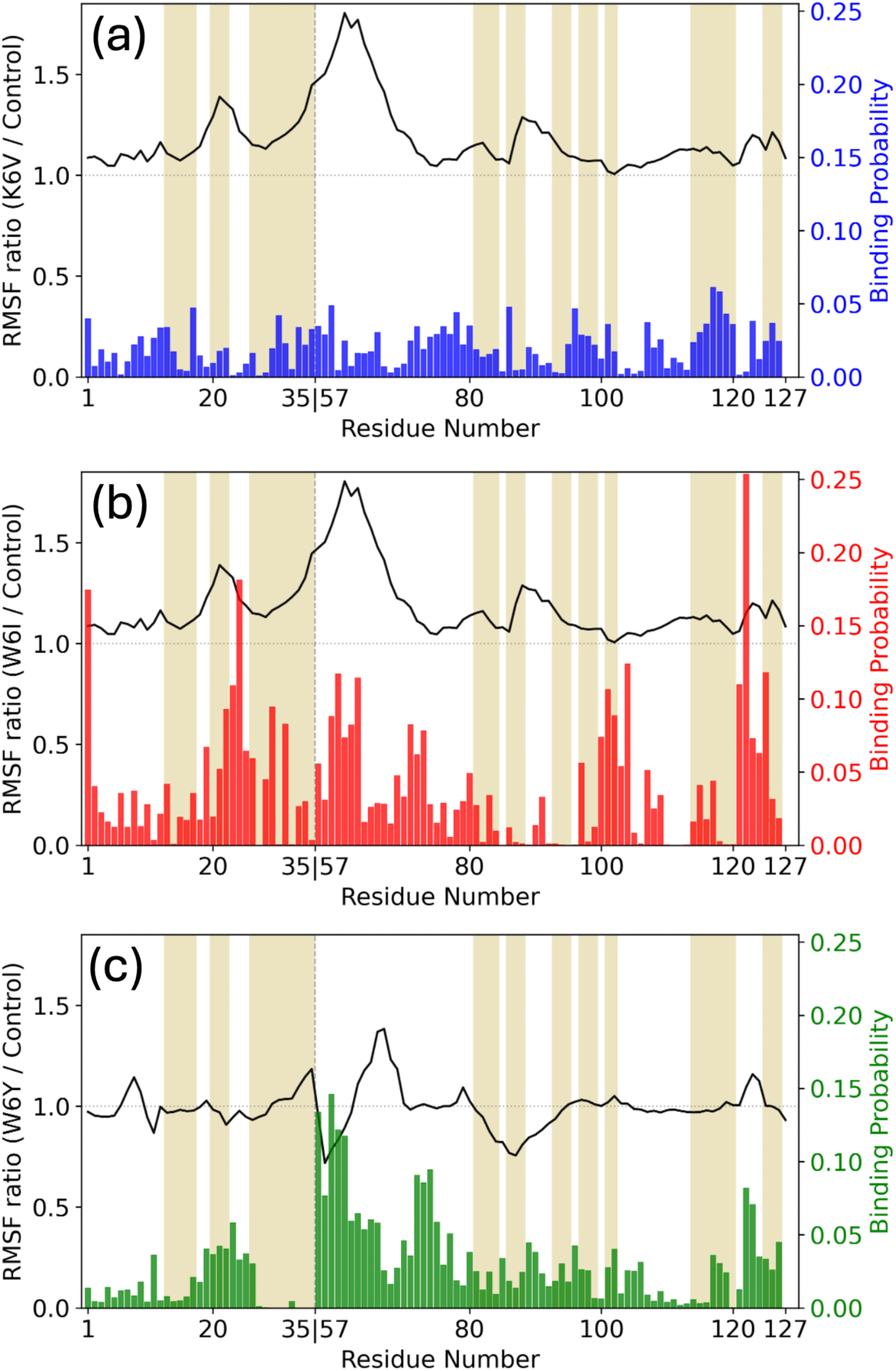
Bar-graphs of the per-residue binding probability of DRI-peptides to the TTR fibril model for (a) DRI-K6V (blue), (b) DRI-W6I (red), and (c) DRI-W6Y (green). The black line marks the ratio of the residue-wise RMSF measured in such simulations divided by the corresponding value in the control simulation. The β-strand regions in the TTR fibril are shaded in yellow.

**Figure 5.**
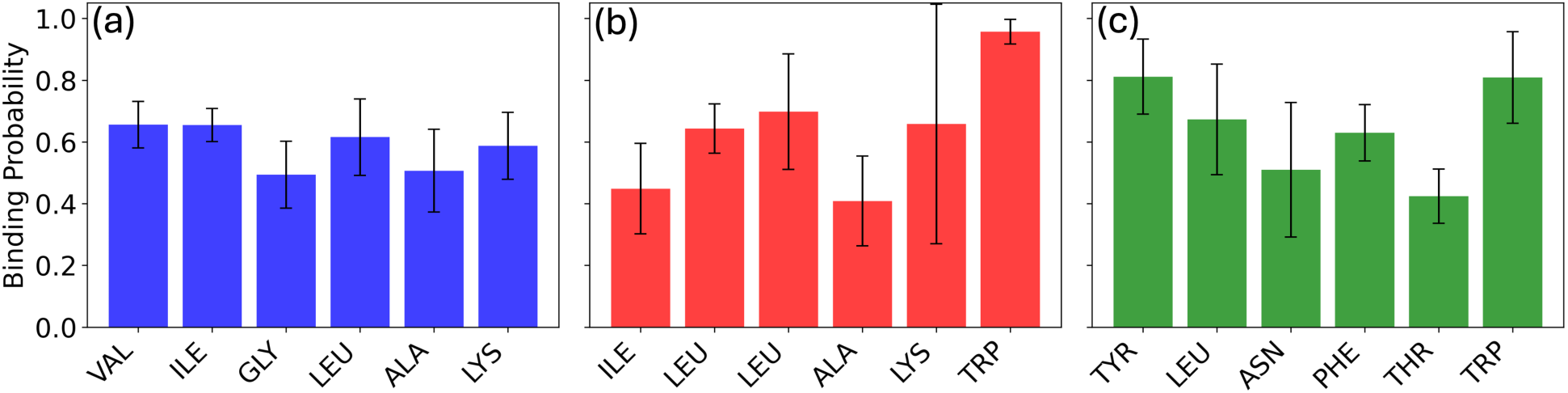
Binding probabilities of the DRI peptide residues binding to the TTR fibril for (a) DRI- K6V (b) DRI-W6I and (c) DRI-W6Y.

We first note that in **Figure 5a** the residues of DRI-K6V have all similar probabilities to bind to the TTR fibril while the distribution is more varied for DRI-W6I (**Figure 5b)** and DRI-W6K (**Figure 5c).** The two peptides include aromatic residues that have binding frequencies of above 90%. Overall, the frequency of DRI-K6V binding with at least one residue to the TTR fibril is with 77% much lower than the corresponding frequencies of DRI-W6I (99%) and DRI-W6Y(95%). The elevated per-residue binding probabilities seen for DRI-W6I and DRI-W6Y are consistent with large aromatic residues that can provide anchoring interactions. Both contain Trp, and DRI- W6Y also contains Phe and Tyr residues which can interact with the TTR fibril surface through π- stacking and extended van der Waals contacts. The more uniform binding probabilities across all DRI-K6V residues suggest a lack of a dominant interaction where contacts are instead distributed across several weak and non-specific interactions.

Using the approach described by Bellaiche and Best^69^ we find that DRI-K6V has a free energy of binding to the fibril of -17 kJ/mol, binding less strongly to the TTR fibril than DRI-W6I (-29.3 kJ/mol) and DRI-W6Y (21.4 kJ/mol). This weaker binding can be also seen in **Figure 4a-4c** where we show the binding probabilities to the TTR residues. As a rule, they are below 7% for DRI-K6V and usually higher for the other two peptides. The distribution for DRI-K6V (**Figure 4a**) is also more even than for DRI-W6I (**Figure 4b**) or DRI-W6Y(**Figure 4c**), and could indicate that DRI- K6V acts through a diffusion mechanism which in other context has been shown to be effective in disrupting specific molecular interactions.^70^ On the other hand, DRI-W6I and DRI-W6Y would have to act through more localized interactions as we find here peaks that correspond to preferred binding sites. In the case of DRI-W6I these correspond to strands C (A25-I26),E (T59-E62), F(E72-I73), I (R104-T106) and L(T118-V122), and for DRI-W6Y to strands E (T59-E62), F(E72- I73) and L(T118-1V22). Hence, both peptides have the polar cavity and the adjacent strand I as preferred binding sites while this is not the cases for DRI-K6V. However, we find for none of the three peptides a signal for significant binding to the region of residues K80-S100, which our control simulations marked as critical.

Note that the initial sites and the binding probabilities differ from what is seen for the inhibitor candidates TabF2 and TabH2 suggested earlier by the Eisenberg group. These peptides were initially designed to bind to the native TTR F and H strands, believed to be involved in the aggregation process, then further optimized for solubility and to limit self-aggregation by the addition of a four-residue arginine tag and an N-methyl group to generate two 16-residue peptides named TabF2 and TabH2.^27, 59^ As described by us in the method section these peptides adopt an extended conformation across the fibril surface (**Figure SF1**), consistent with a surface-capping mechanism. Both N-methylation and positively charged poly-arginine tag optimize solubility and electrostatic recognition at fibril termini but appear to constrain conformational flexibility and restrict penetration into the polar cavity. Notably, despite their high potency as fibril growth inhibitors, TabF2 and TabH2 peptides do not destabilize or degrade existing preformed fibrils.^29^ We expect that this is different for our peptides which clearly interact with the polar cavity and the interior of the fibril, see also the peptide binding probability for the five chains of the TTR fibril model shown in the supplemental **Figure SF2**.

The RMSD plot in **Figure 2** indicates that the effect of this interaction between the peptides and the TTR fibril is for DRI-K6V a loosening of the fibril geometry consistent with the assumption that peptides derived from a protein’s own sequence can bind and inhibit its aggregation.^71^, Visual inspection of the final conformation in the inset of **Figure 2** and of the residue-wise RMSF in **Figure 3** suggests that the locus of the changes is indeed the same as in the high-temperature control simulation, indicating that DRI-K6V exploits the presumed structural weakness of the TTR fibril. Note also that the binding sites correspond to minima in the relative RMSF curves of **Figures 4a-4c** which show the ratio of the RMSF measured in simulations with the peptide present divided by the RMSF values measured in the control simulations. On the other hand, for DRI-W6I and DRI-W6Y we see in **Figure 4b** and **4c** a minimum in the relative RMSF for the K80-S100 segment indicating that presence of the two peptides does not increase the flexibility of that region over that of the control, i.e., does not weaken the interactions between the polar cavity (strand G) and the adjacent strand H. Consistent with our hypothesis are RMSD values in **Figure 2** for the two peptides also lower than for the control, and the final conformations in the inset do not show the loosening of fibril structure seen in the 400K trajectory or in the DRI-K6V simulations.

The above observations are quantified by comparing the time evolution of the radius of gyration (Rg) and solvent accessible surface area (SASA). We list in **Table 2** for these quantities the difference between the averages over three trajectories and the corresponding start values. The absolute values can be found in the Supporting Information **(Table ST3** and **ST4)**.

**Table 2.** Difference between the averages of various quantities, taken over the last 200 ns of three independent trajectories, and the corresponding start values as measured in simulations of the TTR fibril in presence and absence of the three DRI-peptides.

|  | Control | K6V | W6I | W6Y |
| --- | --- | --- | --- | --- |
| **Whole Fibril** |  |  |  |  |
| Rg (Å) | 0.02 (2) | 0.07 (1) | 0.04 (2) | 0.01 (1) |
| Total SASA (Å<sup>2</sup>) | 407 (512) | 1794 (169) | 928 (476) | 699 (267) |
| Hydrophobic SASA (Å<sup>2</sup>) | 692 (278) | 966 (152) | 261 (382) | 158 (260) |
| Hydrophilic SASA (Å<sup>2</sup>) | -285 (294) | 828 (120) | 666 (126) | 541 (116) |
| **N-terminus** |  |  |  |  |
| Rg (nm) | 0.03 (0) | 0.04 (0) | 0.04 (0.01) | 0.03 (0.01) |
| Total SASA (Å<sup>2</sup>) | -894 (279) | -522 (135) | -1047 (140) | -778 (268) |
| Hydrophobic SASA (Å<sup>2</sup>) | -219 (121) | -384 (179) | -566 (109) | -534 (265) |
| Hydrophilic SASA (Å<sup>2</sup>) | -676 (158) | -137 (57) | -481 (31) | -243 (42) |
| **C-terminus** |  |  |  |  |
| Rg (nm) | 0.05 (0.02) | 0.09 (0.01) | 0.07 (0.02) | 0.04 (0.01) |
| Total SASA (Å<sup>2</sup>) | 1301 (417) | 2315 (184) | 1975 (485) | 1476 (207) |
| Hydrophobic SASA (Å<sup>2</sup>) | 911 (287) | 1350 (54) | 828 (408) | 692 (51) |
| Hydrophilic SASA (Å<sup>2</sup>) | 391 (174) | 965 (136) | 1148 (110) | 785 (156) |
| **Cavity** |  |  |  |  |
| Rg (nm) | 0.1 (0.09) | 0.35 (0.20) | 0.16 (0.07) | 0.09 (0.05) |
| Total SASA (Å<sup>2</sup>) | 1032 (485) | 1852 (290) | 1688 (454) | 1291 (183) |
| Hydrophobic SASA (Å<sup>2</sup>) | 897 (301) | 1125 (195) | 981 (307) | 871 (80) |
| Hydrophilic SASA (Å<sup>2</sup>) | 153 (214) | 727 (95) | 705 (163) | 420 (121) |
| **Residues K80-S100** |  |  |  |  |
| Rg (nm) | 0.03 (0.01) | 0.03 (0.02) | 0.01 (0.01) | 0.01 (0.01) |
| Total SASA (Å<sup>2</sup>) | 457 (120) | 511 (265) | 455 (288) | 337 (34) |
| Hydrophobic SASA (Å<sup>2</sup>) | 284 (88) | 440 (197) | 288 (241) | 125 (20) |
| Hydrophilic SASA (Å<sup>2</sup>) | 173 (37) | 71 (69) | 167 (48) | 211 (16) |

The radius of gyration (Rg) is a measure for the compactness of conformations and increases for DRI-K6V and DRI-W6I while changing little in DRI-W6Y and the control. The change is mainly resulting from the C-terminal segment (residues G57-E127) and especially the polar cavity. The loosening of the TTR fibril structure leads for DRI-K6V and DRI-W6I also to a pronounced increase in solvent-accessible surface area (SASA), resulting again mostly from the C-terminal segment and especially the polar cavity. This increase in SASA is mainly resulting from exposure of hydrophilic residues, but in the case of DRI-K6V also from strong exposure of hydrophobic residues.

The changes in Rg and SASA are correlated with a loss of interchain contacts in the C-terminal segment, and again especially pronounced for DRI-K6V and its polar cavity. We list in **Table 3** the difference between the averages of native interchain contacts, that is contacts that exist also in the start conformation, over three trajectories and the corresponding start values. The absolute values can be found in the Supporting Information **Table ST5**. When counting only native contacts, we find in the control an overall loss of 270 (7) of contacts, only slightly larger losses of 304 (15) for DRI-W6I and 283 (12) for DRI-W6Y, but with 360 (7) a much larger loss for DRI- K6V. The loss results mainly from stagging contacts between the layers of the fibril and less from the packing contacts between the N-terminal and C-terminal fragments. For instance, in presence of DRI-K6V 79 more stagging contacts are lost than in the control, but only nine more packing contacts. The loss in stagging contacts is mostly from the cavity where on average 30 more contacts are lost in the DRI-K6V simulation than in the control, and from the segment of residues K80- S100 where in the DRI-K6V simulations 10 more contacts are lost than in the control, while for this segment 11 contacts fewer are lost for DRI-W6I and 21 less for DRI-W6Y.

**Table 3.** Difference between the averages of contacts, taken over the last 200 ns of three independent trajectories, and the corresponding start values as measured in simulations of the TTR fibril in presence and absence of the three DRI-peptides. Only native contacts, that is contacts already found at start, are considered.

|  |  | **Control** | **K6V** | **W6I** | **W6Y** |
| --- | --- | --- | --- | --- | --- |
| **Whole Fibril** | Interchain Contacts | -270 (7) | -360 (7) | -304 (15) | -283 (12) |
|  | Packing contacts | -35 (5) | -46 (2) | -45 (1) | -34 (6) |
|  | Stagging Contacts | -235 (3) | -314 (8) | -259 (11) | -249 (12) |
| **N-terminus** | Stagging Contacts | -50 (2) | -65 (10) | -56 (7) | -60 (5) |
| **C-terminus** | Stagging Contacts | -177 (4) | -232 (9) | -179 (9) | -180 (9) |
| **Cavity** | Stagging Contacts | -92 (6) | -122 (8) | -98 (4) | -109 (7) |
| **Residues K80-S100** | Stagging Contacts | -45 (13) | -55 (13) | -34 (12) | -24 (3) |

Visual inspection of the final conformations shows that the changes to the fibril conformations are mainly in the outer layers which especially in the unphysical 400K simulation, but also in presence of DRI-K6V, get partially separated from the rest of the fibril. For this reason, we have also measured the stacking contacts between chain 1 (the bottom chain) and chain 2, and between chain 4 and chain 5 (the top chain) comparing again simulations where one of the DRI peptides is present with the control simulations. Focusing again on native contacts, we have listed in **Table 4** the differences between averages over the last 200 ns with the corresponding start values. Absolute values are listed in the supplemental **Table ST6**. Quantifying our visual impression of a partial separation of the outer layers, we find that in the control 126 (7) such stagging contacts between the outer chains and the core fibril are lost, but 163 (4) in presence of DRI-K6V and only 135 (14) in presence of DRI-W6I and 127 (12) in presence of DRI-W6Y. While all parts of the TTR fibril loose such stacking contacts, the relative loss is especially strong for the segment K80-S100 where in the control only 6 (about 6%) such contacts are lost, but 12 (12%) in presence of DRI-K6V, 9 (9%) in presence of DRI-W6I and 4 (4%) in presence of DRI-W6Y. Hence, we find that interaction of DRI-K6V with the TTR fibril leads to a partial separation of the outer chains for the segment K80-S100 while this is not the case for the other two peptides. Once started, this separation continues to the polar cavity and finally the full TTR fibril.

**Table 4.** Native stagging contacts between the outer chains and the core of the TTR fibril (i.e. between chain 1 and chain 2, and between chain 4 and chain 5). Shown are the differences between the averages taken over the last 200 ns of three independent trajectories and the corresponding start values as measured in simulations of the TTR fibril in presence and absence of the three DRI- peptides.

|  | **Control** | **K6V** | **W6I** | **W6Y** |
| --- | --- | --- | --- | --- |
| Whole Fibril | -126 (7) | -163 (4) | -135 (14) | -127 (12) |
| N-terminus | -23 (3) | -34 (9) | -25 (2) | -29 (5) |
| C-terminus | -98 (3) | -122 (10) | -100 (14) | -93 (10) |
| Cavity | -52 (5) | -63 (6) | -51 (7) | -58 (5) |
| Residues K80-S100 | -6 (4) | -12 (13) | -9 (9) | -4 (4) |

## CONCLUSIONS

Damages to the heart and other organs in elderly patients are often caused by transthyretin (TTR) amyloid deposits. The lack of treatment options targeting directly these deposits is caused in part due to a lack of a suitable animal model for this TTR amyloidosis making *in vivo* experiments difficult and costly. Hence, detailed knowledge of mechanism by that inhibitors could dissolve the TTR fibril is needed for optimizing potency and selectivity of potential drug candidates prior to experimental validation. This medical question motivated us to look into the possibility to modulate the stability of TTR fibrils with short peptides. Comparing molecular dynamics simulations at an unphysiological high temperature of 400K with such at the physiological temperature of 310K we find indications that disruptions of the fibril structure start with the segment of residues K80-S100 which connects the polar cavity with the adjacent strand H. Using virtual screening we designed three peptides and evaluated in molecular dynamics (MD) simulations at physiological temperatures their ability to destabilize TTR amyloid fibrils. In order to ease future experimental validation, we designed the peptides from D-amino acids as these do not suffer from the short in vivo lifetimes of regular peptide drugs.

In only one case, for DRI-K6V, we see a disruption of the TTR fibril. We find that this process begins with partial segregation of the outer chains starting again with the segment of residues K80- S100 where double as many contacts are lost than in the control. Subsequent losses of stagging contacts in the outer layers in the polar cavity lead to a further weakening that may indicate an avenue for dissolving the TTR fibril. Such weakening of outer β-strand contacts and nearby loops, followed by disruption of interchain networks within the fibril core has been also suggested in other recent work on transthyretin (TTR) fibril destabilization. For instance, it was proposed that the small molecule epigallocatechin gallate (EGCG) destabilizes the mutant V30M TTR fibrils by binding to boundary β-strands and promoting their dissociation.^72^ Additional MD studies suggested that EGCG lowers β-strand content in TTR fibrils, specifically in the cavity region, by destabilizing the L58-I84 contact network at the cavity entrance, while EGC (lacking a gallic acid ester group) binds within the cavity and weakens salt bridges that help maintain its structure.^73^ Unlike this earlier work, our results emphasize the role of the segment K80-S100 as a potential target for modulating TTR fibril stability.

A limitation of our study is that we did not explore potential differences in the efficiency of the peptides to disrupt TTR fibrils between wild type and the mutants causing hereditary forms of TTR amyloidosis as for instance the V30M mutation considered in the computational study^72^ mentioned above. Any medical application of DRI-K6V would have to consider also its binding to competing proteins in the heart or other organs. We hope that by describing a mechanism for disrupting TTR fibrils with DRI-peptides we have built a foundation for targeting these questions in future work.

## DATA AVAILABILITY

The data that support the findings of this study are available in the Supporting Information and will be publicly accessible after publication

## SUPPORTING INFORMATION

The Supporting Information is available free of charge at URL

- Six supplemental tables with statistics of the simulations and supporting data (**TTR_SI.pdf**)

## AUTHOR INFORMATION

### Corresponding Author

Ulrich H. E. Hansmann - Department of Chemistry & Biochemistry, University of Oklahoma, Norman, Oklahoma 73019, United States

### Author

Lucy M. Coleman - Department of Chemistry & Biochemistry, University of Oklahoma, Norman, Oklahoma 73019, United States

### Author Contributions

The manuscript was written through contributions of all authors. All authors have given approval to the final version of the manuscript.

### Notes

The authors have no conflicts to declare.

## ACKNOWLEDGMENTS

The simulations in this work were done using the SCHOONER cluster of the University of Oklahoma, ACCESS resources allocated under grant MCB160005 (National Science Foundation).

## ABBREVIATIONS

ATTR: transthyretin amyloidosis;
ATTRwt: wildtype transthyretin amyloidosis;
ATTRv: hereditary transthyretin amyloidosis;
DRI: D-Retro-Inverso;
FDA: U.S. Food and Drug Administration;
Rg: Radius of gyration;
RMSD: Root mean square deviation;
RMSF: Root mean square fluctuation;
SAA: Serum amyloid A;
SASA: Solvent-accessible surface area;
TTR: transthyretin

