## Supplemental Information for "Modulating Transthyretin Fibril Stability with D-Retro-Inverso Peptides"

Supporting information for this article consists of 6 tables and 2 figures listed on the following pages and two separate files, (**TTR-fibril-coordinates.zip**) a compressed folder with the atomic coordinates of the start and final configurations (as ASCII files in the PDB format) of all trajectories and (**TTR-fibril-topology.zip**) a compressed folder with the topology files and ITP files (separated by chain) of all trajectories of the molecular dynamics simulations considered in this study. The folders contain also README files to help identifying the various files.

### Table of Contents of Supporting Information

**Tables:**

**ST1:** Grouping System based on Properties of Amino Acids. S3

**ST2:** Top 10 peptide candidates ranked by Rosetta binding free energy estimate. ΔG values are reported in Rosetta Energy Units (REU) and represent a computational estimate of binding free energy. S4

**ST3:** Radius of gyration (Rg) measured over the last 200 ns in simulations. Averages over three trajectories for each system are shown, with standard deviations in parenthesis. S5

**ST4:** Solvent accessible surface area (SASA) measured over the last 200 ns in simulations. Averages over three trajectories for each system are shown, with standard deviations in parenthesis. S6

**ST5:** Native contacts measured over the last 200 ns in simulations. Averages over three trajectories for each system are shown, with standard deviations in parenthesis. S7

**ST6:** Native stagging contacts between the outer chains and the core of the TTR fibril (i.e. between chain 1 and chain 2, and between chain 4 and chain 5) measured over the last 200 ns in simulations. Averages over three trajectories for each system are shown, with standard deviations in parenthesis. S8

**Figures:**

**SF1:** Structures of the recently in Ref. 59 proposed inhibitors docked to the TTR fibril: (a) TabF2 in cyan and (b) TabH2 in green. S9

**SF2:** Per-chain binding probability to the TTR fibril for the three considered peptides. S10

**README for additional files in the compressed folder TTR-fibril-coordinates.zip (in separate file)** S11

**README for additional files in the compressed folder TTR-fibril-topology.zip (in separate file)** S13

**Table ST1.** Grouping System based on Properties of Amino Acids

| **Nonpolar, Aliphatic** | **Polar, Uncharged** | **Negatively Charged** | **Aromatic** |
|---|---|---|---|
| Glycine (Gly, G) | Serine (Ser, S) | Aspartate (Asp, D) | Phenylalanine (Phe, F) |
| Alanine (Ala, A) | Threonine (Thr, T) | Glutamate (Glu, E) | Tyrosine (Tyr, Y) |
| Valine (Val, V) | Cysteine (Cys, C) | | Tryptophan (Trp, W) |
| Proline (Pro, P) | Asparagine (Asn, N) | **Positively Charged** | |
| Leucine (Leu, L) | Glutamine (Gln, Q) | Lysine (Lys, K) | |
| Isoleucine (Ile, I) | | Histidine (His, H) | |
| Methionine (Met, M) | | Arginine (Arg, R) | |

**Table ST2.** Top 10 peptide candidates ranked by Rosetta binding free energy estimate. ΔG values are reported in Rosetta Energy Units (REU) and represent a computational estimate of binding free energy.

| Rank | Peptide | ΔG (REU) | Buried SA (Å²) | Interface H-bonds | Unsatisfied H-bonds |
|---|---|---|---|---|---|
| 1 | DRI-KALGIV | -27.135 | 1239.247 | 4 | 3 |
| 2 | DRI-GAAIGW | -27.051 | 1119.133 | 5 | 2 |
| 3 | DRI-YTIGAL | -25.977 | 1357.379 | 8 | 4 |
| 4 | DRI-VALHGF | -25.769 | 1305.09 | 6 | 13 |
| 5 | DRI-STWIFE | -23.989 | 1251.187 | 3 | 12 |
| 6 | DRI-WKALLI | -23.874 | 1438.996 | 3 | 17 |
| 7 | DRI-LKVLDA | -23.752 | 1364.675 | 7 | 12 |
| 8 | DRI-WTFNLY | -23.18 | 1268.806 | 6 | 6 |
| 9 | DRI-GYFLNF | -22.969 | 1343.166 | 4 | 12 |
| 10 | DRI-AVHLFK | -22.804 | 1383.584 | 3 | 5 |

**Table ST3.** Radius of gyration (nm) measured over the last 200 ns in simulations. Averages over three trajectories for each system are shown, with standard deviations in parenthesis.

| | Control | | DRI-K6V | | DRI-W6I | | DRI-W6Y | |
|---|---|---|---|---|---|---|---|---|
| | Initial | Last 200 ns | Initial | Last 200 ns | Initial | Last 200 ns | Initial | Last 200 ns |
| Whole Fibril | 2.36 | 2.38 (0.02) | 2.37 | 2.44 (0.01) | 2.38 | 2.42 (0.02) | 2.37 | 2.38 (0.01) |
| N-terminus | 1.38 | 1.41 (0) | 1.38 | 1.41 (0) | 1.38 | 1.42 (0.01) | 1.38 | 1.41 (0.01) |
| C-terminus | 2.13 | 2.19 (0.02) | 2.15 | 2.24 (0.01) | 2.15 | 2.22 (0.02) | 2.15 | 2.19 (0.01) |
| Cavity | 1.22 | 1.32 (0.09) | 1.23 | 1.6 (0.2) | 1.41 | 1.57 (0.07) | 1.41 | 1.50 (0.05) |
| Residues K80-S100 | 1.33 | 1.36 (0.01) | 1.34 | 1.37 (0.02) | 1.35 | 1.36 (0.01) | 1.35 | 1.36 (0.01) |

**Table ST4.** Solvent accessible surface area (SASA) (Å$^2$) measured over the last 200 ns. Averages over three trajectories for each system are shown, with standard deviations in parenthesis.

| | | **Control** | | **DRI-K6V** | | **DRI-W6I** | | **DRI-W6Y** | |
|---|---|---|---|---|---|---|---|---|---|
| | | Initial | Last 200 ns | Initial | Last 200 ns | Initial | Last 200 ns | Initial | Last 200 ns |
| Total | Whole Fibril | 23108 | 23515 (512) | 23034 | 24828 (169) | 22876 | 23804 (476) | 22750 | 23449 (267) |
| | N-terminus | 9127 | 8233 (279) | 8706 | 8184 (135) | 8677 | 7630 (140) | 8685 | 7907 (268) |
| | C-terminus | 13981 | 15282 (417) | 14328 | 16643 (184) | 14199 | 16174 (485) | 14066 | 15542 (207) |
| | Cavity | 4257 | 5289 (485) | 4425 | 6277 (290) | 4442 | 6130 (454) | 4305 | 5596 (183) |
| | Residues K80-S100 | 4421 | 4878 (120) | 4717 | 5228 (265) | 4706 | 5161 (288) | 4610 | 4947 (34) |
| Hydrophobic | Whole Fibril | 11330 | 12022 (278) | 11778 | 12744 (152) | 11765 | 12026 (382) | 11774 | 11932 (260) |
| | N-terminus | 4626 | 4407 (121) | 4721 | 4337 (179) | 4561 | 3995 (109) | 4671 | 4137 (265) |
| | C-terminus | 6704 | 7615 (287) | 7056 | 8406 (54) | 7204 | 8032 (408) | 7104 | 7796 (51) |
| | Cavity | 1713 | 2610 (301) | 1946 | 3071 (195) | 2115 | 3096 (307) | 2003 | 2874 (80) |
| | Residues K80-S100 | 2084 | 2368 (88) | 2211 | 2651 (197) | 2228 | 2516 (241) | 2220 | 2345 (20) |
| Hydrophilic | Whole Fibril | 11778 | 11493 (294) | 11256 | 12084 (120) | 11111 | 11777 (126) | 10976 | 11517 (116) |
| | N-terminus | 4501 | 3825 (158) | 3984 | 3847 (57) | 4116 | 3635 (31) | 4014 | 3771 (42) |
| | C-terminus | 7277 | 7668 (174) | 7272 | 8237 (136) | 6995 | 8143 (110) | 6962 | 7747 (156) |
| | Cavity | 2526 | 2679 (214) | 2479 | 3206 (95) | 2328 | 3033 (163) | 2302 | 2722 (121) |
| | Residues K80-S100 | 2337 | 2510 (37) | 2506 | 2577 (69) | 2478 | 2645 (48) | 2390 | 2601 (16) |

**Table ST5.** Native contacts measured over the last 200 ns in simulations. Averages over three trajectories for each system are shown, with standard deviations in parenthesis.

| | | **Control** | | **DRI-K6V** | | **DRI-W6I** | | **DRI-W6Y** | |
|---|---|---|---|---|---|---|---|---|---|
| | | Initial | Last 200 ns | Initial | Last 200 ns | Initial | Last 200 ns | Initial | Last 200 ns |
| Native Interchain Contacts | Whole Fibril | 1190 | 920 (7) | 1229 | 869 (7) | 1199 | 895 (11) | 1208 | 925 (17) |
| Native Packing Contacts | | 126 | 91 (5) | 128 | 82 (2) | 123 | 78 (1) | 125 | 91 (6) |
| Native Stagging Contacts | | 1064 | 829 (3) | 1101 | 787 (8) | 1076 | 817 (11) | 1083 | 834 (12) |
| Native Stagging Contacts | N-terminus | 262 | 212 (2) | 269 | 204 (10) | 270 | 214 (7) | 266 | 206 (5) |
| | C-terminus | 747 | 570 (4) | 775 | 543 (9) | 749 | 570 (9) | 765 | 585 (9) |
| | Cavity | 263 | 171 (6) | 280 | 158 (8) | 270 | 172 (4) | 286 | 177 (7) |
| | Residues K80-S100 | 194 | 149 (13) | 195 | 140 (13) | 188 | 154 (12) | 192 | 168 (3) |

**Table ST6.** Native stagging contacts between the outer chains and the core of the TTR fibril (i.e. between chain 1 and chain 2, and between chain 4 and chain 5) measured over the last 200 ns in simulations. Averages over three trajectories for each system are shown, with standard deviations in parenthesis.

| | **Control** | | **DRI-K6V** | | **DRI-W6I** | | **DRI-W6Y** | |
|---|---|---|---|---|---|---|---|---|
| | Initial | Last 200 ns | Initial | Last 200 ns | Initial | Last 200 ns | Initial | Last 200 ns |
| Whole Fibril | 530 | 404 (7) | 543 | 380 (4) | 538 | 403 (14) | 530 | 403 (12) |
| N-terminus | 125 | 102 (3) | 132 | 98 (9) | 132 | 107 (2) | 128 | 99 (5) |
| C-terminus | 377 | 279 (3) | 383 | 261 (10) | 381 | 281 (14) | 376 | 283 (10) |
| Cavity | 135 | 83 (5) | 139 | 76 (6) | 136 | 85 (7) | 143 | 85 (5) |
| Residues K80-S100 | 96 | 90 (4) | 97 | 85 (13) | 95 | 86 (9) | 94 | 90 (4) |

**Figure SF1.** Structures of the recently in Ref. 59 proposed inhibitor docked to the TTR fibril: (a) TabF2 (in cyan) and (b) TabH2 (in green).

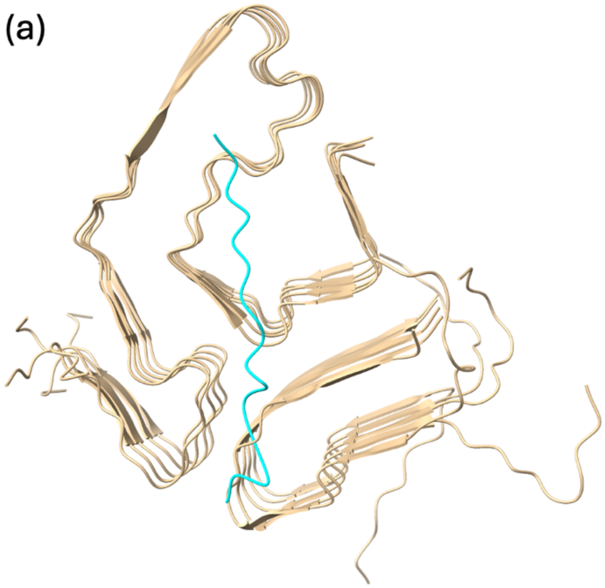


Saelices, L.; Chung, K.; Lee, J. H.; Cohn, W.; Whitelegge, J. P.; Benson, M. D.; Eisenberg, D. S. Amyloid seeding of transthyretin by ex vivo cardiac fibrils and its inhibition. *Proc Natl Acad Sci U S A* **2018**, *115* (29), E6741-E6750. DOI: 10.1073/pnas.1805131115.

**Figure SF2.** Per-chain binding probability to the TTR fibril for the three considered peptides over the last 200 ns.

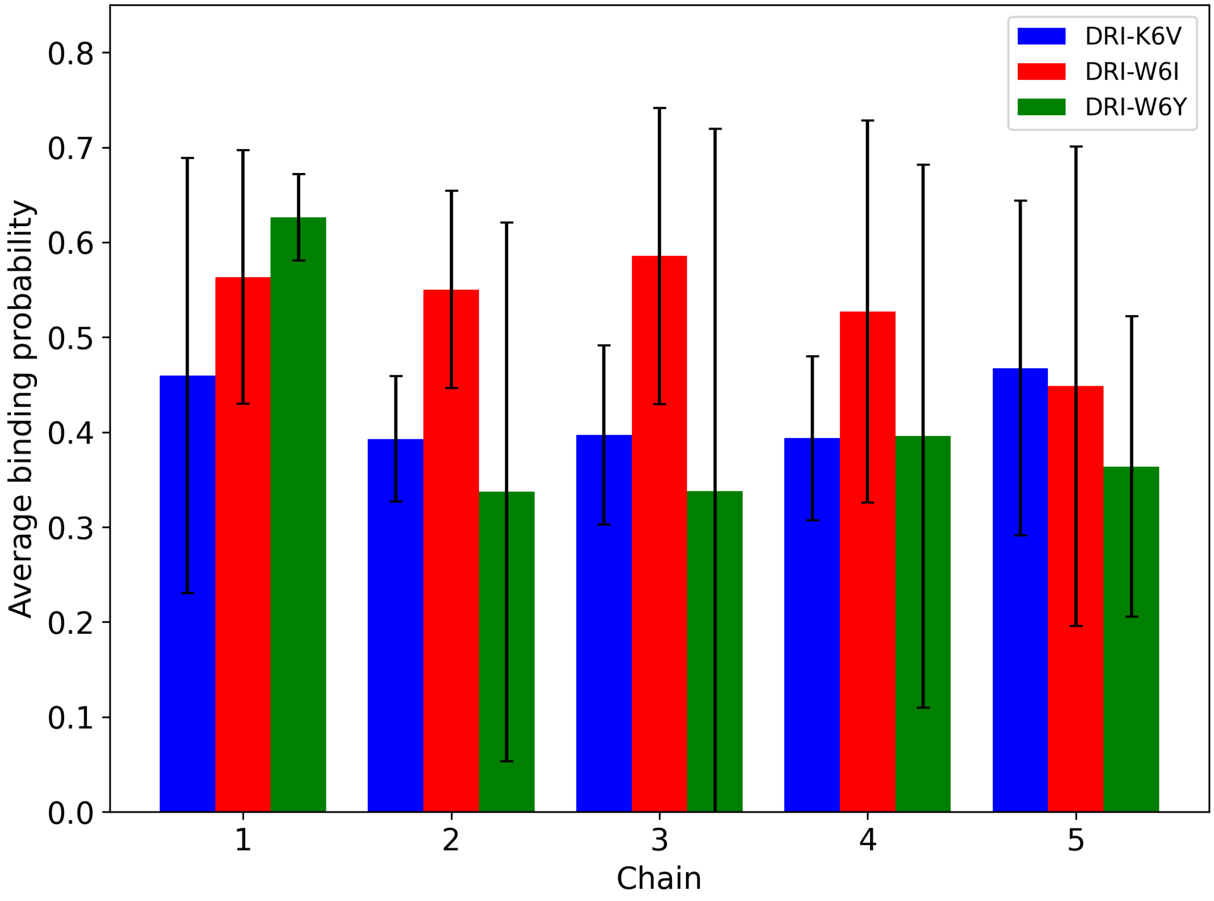
